# Herpes Simplex Virus Mistyping due to HSV-1 x HSV-2 Interspecies Recombination in Viral Gene Encoding Glycoprotein B

**DOI:** 10.1101/2020.02.25.965426

**Authors:** Amanda M. Casto, Meei-Li Huang, Hong Xie, Keith R. Jerome, Anna Wald, Christine Johnston, Alexander L. Greninger

## Abstract

Human herpes simplex viruses (HSV) 1 and 2 are most often typed via molecular assays. Here we describe the first known case of HSV mistyping due to a previously undescribed HSV-1 x HSV-2 recombination event in UL27, the gene that encodes glycoprotein B. This is the first reported HSV interspecies recombination event impacting this gene, which is frequently used as a target for diagnostics and experimental therapeutics.

## Introduction

Human herpes simplex viruses (HSV) 1 and 2 cause chronic, incurable infections in 3 billion and 500 million people worldwide, respectively [1,2]. HSV-1 typically causes orofacial lesions while HSV-2 usually causes genital ulcer disease. However, there is extensive overlap in the spectrum of disease that can be attributed to each virus. The identification of the species of HSV responsible for an infection can provide important diagnostic and prognostic information. The 2015 CDC STD Treatment Guidelines recommend distinguishing between HSV-1 and HSV-2 as the cause of genital herpes because infections due to HSV-1 have a much lower rate of recurrence [3]. IDSA Guidelines also recommend the use of an HSV typing assay in the diagnosis of CNS infections as HSV-1 tends to cause a severe encephalitis with high rates of mortality while HSV-2 causes a milder but recurrent meningitis [4].

Molecular HSV typing is performed either using commercial test kits or testing assays developed by clinical laboratories. All fourteen FDA-cleared HSV typing assays distinguish between HSV-1 and HSV-2 using species-specific probes that recognize genomic DNA or mRNA sequences. As a result, naturally occurring genetic variation in HSV can impact the results of typing assays.

## Methods

### Sample, Typing, and Sequencing Analysis

The sample described in this article, CT_Sample9, was collected in the course of a clinical trial. All trial participants consented to the genetic analysis of their HSV samples.

HSV samples from all trial participants were sent to the University of Washington Clinical Virology Laboratory for typing using a PCR based assay that amplifies a portion of the UL27 gene with a different sequence in HSV-1 and HSV-2 [5]. To identify antiviral resistance mutations, species-specific primers were then used to sequence the UL23 (encoding the HSV thymidine kinase) and UL30 (encoding the HSV DNA polymerase) genes.

CT_Sample9 was additionally subjected to whole genome sequencing using a hybridization probe capture technique developed specifically for HSV [6]. A CT_Sample9 consensus sequence for the unique regions of the genome (unique long and unique short, which comprise 78% of the genome) was created via *de novo* assembly of sequencing reads in Geneious v10 [7].

### Phylogenetics and Recombination Detection

The CT_Sample9 consensus sequence was aligned to HSV-1 Strain 17 (NC_001806.2) and HSV-2 Strain SD90e (KF781518.1) using MAFFT [8]. The portion of this alignment encoding UL27 was scanned for recombination using BootScan as implemented in the recombination detection program (RDP) [9].

## Results

### Conflicting assay results observed for CT_Sample9

The HSV typing assay used by our clinical laboratory [5] indicated that sample CT_Sample9 was positive for HSV-1. To assess for antiviral resistance, we attempted to sequence the UL23 and UL30 genes using HSV-1 specific primers. However, the genes failed to amplify. A subsequent attempt to amplify these genes using HSV-2 specific primers was successful.

### Genome Consensus Sequence for CT_Sample9 consistent with HSV-2

We next performed whole genome sequencing on CT_Sample9. *De novo* assembly of sequencing reads resulted in a 108,998 bp contig and a 16,668 bp contig at an average read depth of more than 2,000x that aligned to the unique long and unique short regions, respectively, of both HSV-1 strain 17 and HSV-2 strain SD90e. The CT_Sample9 consensus sequence differed from Strain 17 by 21,829 single nucleotide changes and from SD90e by only 287 changes, demonstrating that CT_Sample9 is HSV-2 and not HSV-1.

### HSV-1 x HSV-2 Recombination Event observed in Glycoprotein B in CT_Sample9

We reviewed the UL27 gene in an alignment of the HSV-1 strain 17, HSV-2 SD90e, and the CT_Sample9 consensus genomes. A 17 bp portion of this gene used in the HSV typing assay suggested that CT_Sample9 was an HSV-1 sample [5]. The primers for this assay aligned to sequences that were identical in HSV-1, HSV-2, and CT_Sample9 (Figure 1A). However, the probe region differs between the HSV-1 and HSV-2 reference by five single nucleotide changes. We noted that the CT_Sample9 consensus genome was identical to the HSV-1 reference sequence in the probe region. Outside the probe region, there were three additional nucleotide positions upstream and five additional nucleotide positions downstream where the HSV-1 and HSV-2 references differed and the CT_Sample9 sequence matched the HSV-1 reference sequence. Two of these changes were non-synonymous (I527V and V529I). Outside of the region containing these 13 loci, the CT_Sample9 sequence matched the HSV-2 reference at all loci in UL27 where the HSV-1 and HSV-2 references differed. This suggested that the CT_Sample9 genome contained an HSV-1 recombination event of at least 114 bp (maximum length 177 bp) within UL27 which spanned these 13 loci. The recombinant region had over 2,000-fold read coverage and none of the alleles defining the recombination event were present in less than 98.0% of reads.

**Figure 1:**
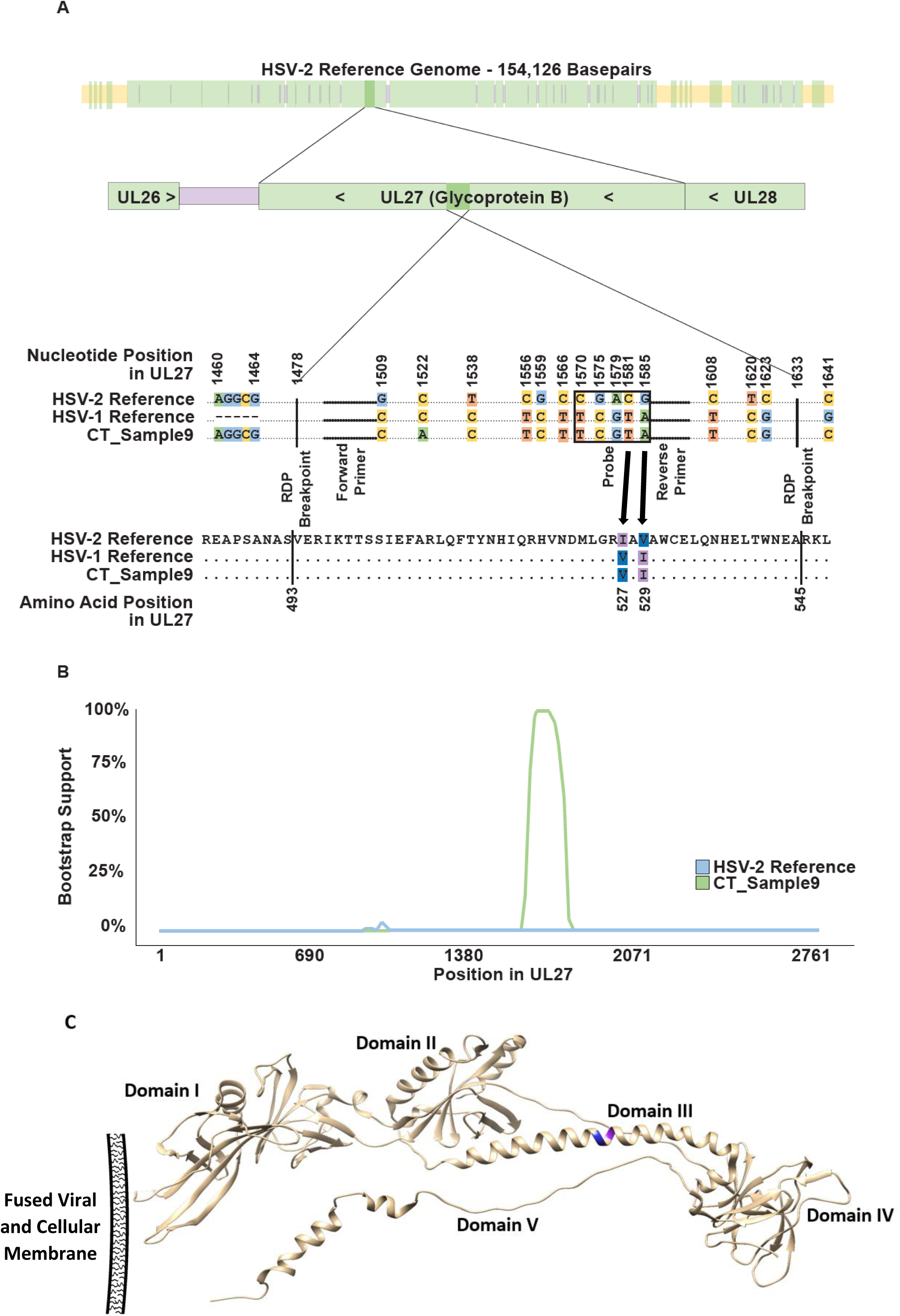
A) Nucleotides involved in recombination event in UL27. The top bar shows the location of the UL27 gene (shown in dark colors) within the HSV-2 genome (faded out). Genic regions in the HSV-2 genome are shown in green while intragenic sequence outside the terminal and internal repeat regions is in purple. Intragenic regions within the terminal and internal repeat regions are in yellow. The second bar shows UL27 in its genic neighborhood with the recombinant region highlighted. UL27 is in the reverse orientation in the HSV genome. In the alignment, nucleotides that are identical in all sequences are represented by dots while nucleotides that differ among sequences are represented by letters. Dots shown in bold represent the locations of the forward and reverse primers as labeled. The probe region is boxed. Recombination event breakpoints as identified by RDP are labeled. Non-synonymous differences between HSV-1 and HSV-2 are indicated with arrows pointing to corresponding amino acid changes. B) BootScan plot of the HSV-2 reference and CT_Sample9 sequences for UL27. The HSV-1 reference sequence is used as the query sequence. C) Postfusion protein conformation of glycoprotein B of HSV-1. Residue 527 (V) is shown in blue and residue 529 (I) is shown in purple. The protein is divided in five structural domains. Residues 527 and 529 are within Domain III of glycoprotein B, which is thought to be important in the binding of three glycoprotein peptides together to form the active trimer. The location of the fused viral and cellular membranes relative to the different domains of the postfusion protein is indicated.

We also used the program Bootscan to examine UL27 in CT_Sample9 for evidence of recombination. The BootScan plot is shown in Figure 1B. The p-value for the recombination event as detected by BootScan is 1.642 × 10^−17^.

### Single Nucleotide Variants also found in primer and probe sequences for typing assay

Due to the recombination event described above, CT_Sample9 was typed as HSV-1 though genome-wide evidence suggested the sample is HSV-2. As the typing assay relies on the homology of its primers and probes with a viral genome, genetic variation other than HSV-1 x HSV-2 recombination in the genomic region targeted by the assay also had the potential to interfere with the assay’s results. We scanned 229 publicly available HSV-1 UL27 sequences and found that two (0.9%) had at least one single nucleotide change in the assay’s forward primer, four (1.8%) had at least one change in the reverse primer, and two (0.9%) had a change in the probe sequence. Seven of the eight HSV-1 samples with these changes were collected in Seattle. The eighth was collected in Kenya. Four HSV-2 genomes out of 459 publicly available genomes differed from the assay’s HSV-2 probe sequence by the same single nucleotide change. All were collected in Seattle. There were no HSV-2 genomes that differed from either of the primer sequences.

## Discussion

This case illustrates how the reliability of DNA/RNA sequence-based diagnostics for infectious diseases is determined by our knowledge of pathogen genomic variation. Both the recombination event in UL27 and the single nucleotide variants that we identified within the primer and probe regions are present at low frequencies among sequenced HSV-1 and HSV-2 samples. However, sequenced samples poorly represent global HSV-1 and HSV-2 populations, having been collected from relatively few geographic locations [10]. As exceedingly few of the thousands of HSV specimens that are typed for clinical purposes every year are subsequently sequenced, it is difficult to predict how often mistyping or suboptimal assay performance occurs due to HSV genomic variation. There are many different HSV typing assays that all target different genomic regions and the primer and probe sequences for commercial assays are not publicly available, further compounding this uncertainty. Of the 14 FDA-approved commercial HSV typing assays [11], information on the target gene(s) is available for eleven. Seven out of these 11 assays target at least one HSV gene where an HSV-1 x HSV-2 recombination event has been described.

This is the first described instance of an HSV-1 x HSV-2 recombination event affecting one of the HSV genes that encodes a glycoprotein. UL27 is a well-studied, essential gene that encodes a protein (Glycoprotein B) critical for viral cell entry. The gene is commonly used as a target for diagnostics and therapeutics, including vaccine candidates. It contains 27 described immune epitopes in HSV-2, one of which is interrupted by the recombination event [12]. As there is no described epitope in the same region in HSV-1, this raises the possibility that the CT_Sample9 recombination event evolved as a means of immunological escape.

In summary, we have described an HSV-1 x HSV-2 recombination event resulting in HSV species mistyping. This is the first reported HSV interspecies recombination event in UL27, a gene of diagnostic and therapeutic importance. This case demonstrates that our knowledge of HSV genomic variation remains limited and shows how these limitations can impact the performance of clinical diagnostic tests.

## References

1. Looker KJ, Magaret AS, May MT, et al. Global and Regional Estimates of Prevalent and Incident Herpes Simplex Virus Type 1 Infections in 2012. PLoS ONE. 2015; 10(10):e0140765.

2. Looker KJ, Magaret AS, Turner KME, Vickerman P, Gottlieb SL, Newman LM. Global estimates of prevalent and incident herpes simplex virus type 2 infections in 2012. PLoS ONE. 2015; 10(1):e114989.

3. 2015 STD Treatment Guidelines [Internet]. 2019 [cited 2019 Aug 13]. Available from: https://www.cdc.gov/std/tg2015/default.htm

4. Miller JM, Binnicker MJ, Campbell S, et al. A Guide to Utilization of the Microbiology Laboratory for Diagnosis of Infectious Diseases: 2018 Update by the Infectious Diseases Society of America and the American Society for Microbiologya. Clinical Infectious Diseases. 2018; 67(6):e1–e94.

5. Corey L, Huang M-L, Selke S, Wald A. Differentiation of herpes simplex virus types 1 and 2 in clinical samples by a real-time taqman PCR assay. J Med Virol. 2005; 76(3):350–355.

6. Greninger AL, Roychoudhury P, Xie H, et al. Ultrasensitive Capture of Human Herpes Simplex Virus Genomes Directly from Clinical Samples Reveals Extraordinarily Limited Evolution in Cell Culture. mSphere. 2018; 3(3).

7. Kearse M, Moir R, Wilson A, et al. Geneious Basic: an integrated and extendable desktop software platform for the organization and analysis of sequence data. Bioinformatics. 2012; 28(12):1647–1649.

8. Katoh K, Standley DM. MAFFT multiple sequence alignment software version 7: improvements in performance and usability. Mol Biol Evol. 2013; 30(4):772–780.

9. Martin DP, Murrell B, Golden M, Khoosal A, Muhire B. RDP4: Detection and analysis of recombination patterns in virus genomes. Virus Evol. 2015; 1(1):vev003.

10. Casto AM, Roychoudhury P, Xie H, et al. Large, stable, contemporary interspecies recombination events in circulating human herpes simplex viruses. J Infect Dis. 2019;.

11. US Food and Drug Administration, Nucleic Acid Based Tests, List of Microbial Tests, Herpes Simplex Viruses [Internet]. US Food and Drug Administration; 2020 [cited 2020 Feb 24]. Available from: http://www.fda.gov/medical-devices/vitro-diagnostics/nucleic-acid-based-tests

12. Laing KJ, Magaret AS, Mueller DE, et al. Diversity in CD8(+) T cell function and epitope breadth among persons with genital herpes. J Clin Immunol. 2010; 30(5):703–722.

